# MILD TRAUMATIC BRAIN INJURY IMPAIRS EPISODIC MEMORY IN RATS

**DOI:** 10.1101/2025.04.28.651041

**Authors:** Gabriel D. Nah, Mira N. Antonopoulos, Andrea G. Hohmann, Nicholas Port, Jonathon D. Crystal

## Abstract

Mild traumatic brain injury (mTBI) is the most common type of traumatic brain injury. Symptoms following mTBI fall into physical, emotional, sleep, and cognitive categories, with memory deficits being a commonly documented sequelae. Whereas many animal models of mTBI exist, relatively few studies have examined the cognitive deficits of mTBI with human-like cognitive tasks. The Wayne State University Closed Head Weight Drop Model recapitulates critical physical elements of sport-related concussions and trauma-based mTBI. However, until now, this model has not previously been evaluated using a human-like memory task. Rats were trained in an odor-based item-in-context task that dissociates episodic and non-episodic memory (Panoz-Brown et al., Current Biology, 2016). The animals then underwent either a weight drop or a sham procedure. After the manipulation, animals were assessed in the item-in-context task. Episodic memory was significantly impaired in the injured rats by over 10% but not in the sham rats. Non-episodic memory was not impaired in either group. Additionally, a time-course immunohistochemical analysis of the hippocampus was performed to examine possible time-dependent changes in ionized calcium-binding adaptor molecule 1 (iba1), a marker of activated microglia/macrophages and glial fibrillary acidic protein (GFAP), a marker of astrocytes. Concussion injury was associated with time-dependent morphological changes in astrocytes and microglia in injured rats compared to sham rats. This study is the first to document episodic memory impairment in an animal model of mTBI.

## INTRODUCTION

Mild traumatic brain injury (mTBI), or concussion, is the most common type of traumatic brain injury, and it affects over two million people each year. Among the various symptoms following mTBI, memory impairment is among the most common (Masse et al., 2019; Qin et al., 2018). Memory is an important aspect of cognition. The ability to encode, store, and retrieve information is a critical aspect of memory. Developing animal models of human memory disorders requires two components: a model of the biological disorder and a sensitive measure of behavior that taps into the types of impairments associated with the disorder. Failures to translate from preclinical models to clinical conditions in people may stem from deficiencies in either component. Without a working model for both components, researchers cannot thoroughly examine the extent of damage related to mTBI in humans (Hiskens, Angoa-Perez, Schneiders, Vella, & Fenning, 2019). The present work focuses on developing an animal model of mTBI and an impairment of episodic memory.

Currently, limited treatment options exist for mTBI. The recommended therapy involves rest and reduced activity until the symptoms have resolved (CDC, 2019). However, this therapy is only intended to manage the acute symptoms, and currently, no drugs exist to prevent the long-term effects of TBI (Dewitt, Perez-Polo, Hulsebosch, Dash, & Robertson, 2013). Investigating better therapeutic options for TBI is an important objective, and developing a valid animal model of sports-related mTBI is an initial step in evaluating therapeutic drugs in animal studies.

Although numerous rodent models of mTBI have been developed (e.g., fluid percussion injury, controlled cortical impact, etc.), none of these models have been validated to show human-like behavioral deficits (Romeu-Mejia, Giza, & Goldman, 2019). Efforts to model mTBI in nonhumans are often characterized by injuries that do not mimic the biomechanics of sport-related injuries in people (Hiskens et al., 2019). Therefore, a significant gap in the literature is the lack of an animal model of concussion that both recapitulates the biomechanics of sport-related injuries and the types of impairments to complex aspects of cognition observed in human concussion. The Wayne State University Closed-Head Weight drop model (WDM) can produce an mTBI with similar biomechanics to sport-related injuries in humans without the limitations other models have (i.e., surgery, skull fractures, high mortality rate, restrained animal) (Masse et al., 2019; Qin et al., 2018).

The hippocampus is key in the formation of new memories and memory storage (Parkin, 1996). One type of hippocampal-dependent memory is episodic memory (Tulving & Markowitsch, 1998). Episodic memory can be described as the ability to remember specific events or experiences in sequential order and the places they occur (Crystal, 2021; Eichenbaum, 2001; Tulving, 2002). Episodic memory can be observed in nonhuman animals (Crystal, 2009; Dere, Kart-Teke, Huston, & De Souza Silva, 2006; Morris, 2001). Studies have shown that rats can demonstrate episodic memory functions similar to humans. Panoz-Brown and colleagues (2016) developed an item-in-context task to demonstrate that rats can use episodic memory while differentiating from non-episodic memory solutions. This task was selected as the memory assessment for this study because it demonstrates that rats can model a key aspect of human memory (Crystal, 2021; Morris, Weeden, Churchwell, & Kesner, 2013).

Studies show hippocampal dysfunction can impair episodic memory function (Bai et al., 2009; La et al., 2019; Mormino et al., 2009; Panoz-Brown et al., 2018). MTBI may impair memory by damaging the hippocampus. The hippocampus is responsible for binding spatial and non-spatial information, creating context-dependent memories, a defining feature of episodic memory (Lee & Jung, 2017). Lesions in parts of the hippocampus can impede the ability to form associations between odors and contexts (Morris, 2001). Consequently, when the hippocampus is damaged during an mTBI, it may impair episodic memory. Researchers use markers of injury and inflammation to measure damage in the brain. Activation of astrocytes and microglia are used to assess injury and inflammation. Astrocytes are usually involved in homeostasis and blood flow to the central nervous system. During brain injury, astrocytes experience increased activation, called astrogliosis (Sofroniew & Vinters, 2010; Zhou et al., 2020). Microglia also exhibit a similar response and play a role in tissue regeneration. During brain injury, microglia become activated and secrete inflammatory cytokines (Grovola et al., 2021; Lafrenaye, Todani, Walker, & Povlishock, 2015; Zhou et al., 2020). Those inflammatory responses suggest that astrocytes and microglia can be valuable markers of injury and inflammation in the brain following mTBI.

The aim of this study was divided into two parts. First, we tested whether this WDM induces episodic memory impairment in rats. We used the episodic memory task described by Panoz-Brown et al., (2016) and tested episodic memory function before and after weight drop or sham procedures. We hypothesized that rats with an mTBI, but not sham rats, would show a decline in episodic memory performance. Second, we measured the morphology of astrocytes and microglia to quantify hippocampal damage following mTBI in rats. We hypothesized that rats with an mTBI would show morphological changes in astrocytes and microglia.

## METHODS

### Animals

Male and female Sprague-Dawley rats (Harlan, Indianapolis, IN) were housed individually and maintained on a 12-hour light/dark cycle with light onset at 7:50 and offset at 19:50 EST. 12 male and female Sprague-Dawley rats were used for behavioral assessments. Thirty male and female Sprague-Dawley rats were used for immunohistochemical analysis. Upon arrival in the lab at the age of 7 weeks, the rats were given unlimited access to 5012-Rat-Diet (PMI Nutrition International, St. Louis, MO) for one week. Next, male rats received 15 g/day, and female rats received 11g/day of food if selected for behavioral testing. Water was available ad libitum, except during brief testing sessions. All procedures followed national guidelines and were approved by the Bloomington institutional animal care and use committee at Indiana University. One female rat was euthanized at the beginning of the study due to health complications.

### Apparatus

Two open-field arenas were used for odor presentation and served as distinctive contexts. Arena-A was circular in shape with a 94-cm diameter white floor and enclosed with a 30-cm high wall constructed from white acrylic plexiglass. Eighteen circular holes (5 cm diameter, 2.5 cm deep) were arranged in two concentric circles with 6 and 12 holes in the inner and outer rings, respectively. Arena-B was circular in shape with a 46-cm diameter floor and a transparent 30-cm high wall. The floor of Arena-B contained an array of 3 concentric circles that alternated in color. The innermost circle was black, the middle circle was white, and the outer circle was black. The inside of Arena-B consisted of eight equidistant circular holes (5 cm diameter, 2.5 cm deep) positioned along the wall. Each condiment cup (59 ml) was firmly snapped inside a hole to lay flush with the floor and was covered with a plastic lid loosely placed on top. White noise was used throughout to mask outside noise. After each animal’s daily session, both arenas were cleaned with a 2% chlorhexidine solution.

### Description of Odors

Odors were presented using opaque plastic lids. Approximately 40 lids were stored in a sealed container containing a spice or oil odorant to odorize the lids. The lids were stored for at least two weeks before odor presentation began. The odors included almond oil, amaretto oil, banana, asparagus, blueberry oil, brandy oil, butterscotch oil, caraway seed, celery seed, chicory root, cinnamon, coffee oil, cumin, dill weed, garlic powder, hickory smoke, honey oil, horseradish, Irish cream oil, lavender, lemon zest, maple, menthol-eucalyptus, Mexican oregano, mustard seed, onion powder, orange oil, pecan oil, pineapple oil, root beer oil, rosemary leaf, sage leaf, sesame oil, spinach powder, strawberry oil, summer savory, thyme, tomato, Mexican vanilla, and watermelon oil.

### Episodic Memory Task

Episodic memory was assessed in the Item-in-Context task using a list of 16 odors described in Panoz-Brown et al. (2016). Rats were tested once daily, five to seven days weekly, for 12 weeks. Between trials, the rat was placed in a cage identical to its home cage, but it did not contain food, water, or bedding. For each session, the order of the contexts, lists of odors, and location of odors within each arena were randomly selected. Each correct choice during the task was rewarded with a single chocolate pellet (F0229, Bio-Serv, Frenchtown, NJ). The rats progressed through pretraining, approximately ten days of odor span training, five days of preliminary training, ten days of the memory task with two context transitions, and 15 days of the memory task with three context transitions (Panoz-Brown et al., 2016). A new plastic lid (carrying the scheduled odor) was used during each odor presentation to preclude any role of rat-induced scent marking.

### Pretraining

Pre-training involved acclimating the rats to the arenas and the task. First, cups baited with one chocolate pellet were placed in every location throughout one arena, and the rat was placed in the arena and permitted to forage until all the chocolate was consumed. This was immediately followed by repeating the same procedure in the other arena. Once the rat consumed all pellets within five minutes, it began lid shaping. In a lid-shaping session, the rat underwent 15 trials in the first arena and 15 in the second arena. In each trial, one baited cup was placed randomly in the arena. Initially, an unscented lid was placed near the cup. As the rat progressed through the training, the lid was gradually moved on successive trials until it entirely covered the baited cup. Rats moved on to odor span training when they consistently displaced the lid from 100% lid coverage and consumed the food.

### Odor Span Training

Sixteen odors were randomly selected for each session and presented in a different order in each arena. In the first trial, one baited cup covered with one odorized lid was placed randomly in the first arena. Then, the rat was placed in the arena and allowed to find the odor, displace the lid, and consume the food. The first (old) and second (new) odors were placed at randomly selected locations in the subsequent trial. The rat was then given a choice between the old, unbaited odor (S-) and the new, baited odor (S+). A correct choice was defined as displacing the lid with the S+odor, and an incorrect choice was defined as displacing the lid with the S-odor. A lid displacement was defined as a vertical or horizontal movement of the lid. When an incorrect choice occurred, the rat was allowed to search the arena until the correct choice was made, but only the first choice was included in the data analysis.

As the session progressed, the previous S-odors were simultaneously presented in the arena, so each correct response led to a subsequent increase in the number of S-odors. This continued until the list was completed or until an incorrect choice was made. After an incorrect choice, all of the accumulated odors were removed for the remainder of the session in the arena, and the task resumed, with the subsequent odor on the list being presented as the sole lid. The number of presented odors continued to increase with each correct choice. After the list was completed in the first arena, the same procedure was completed in the second arena, using the same list of odors in a different random order. All 11 items were presented as new in the second arena using a new randomly selected order and locations.

### Preliminary Training

Preliminary training involved a two-alternative forced-choice task. It consisted of 32 trials, with 16 trials in each arena. It was similar to the odor span task, but every trial after the first trial consisted of only two odors, one new (S+) and one old (S-). The S-odor for each trial was randomly chosen from the odors previously presented in that arena. The task proceeded until all 16 odors were presented in the first arena. Then, the procedure was repeated in the second arena using the same odors presented in a new random order. Locations of lids were randomly selected.

### Context Transition Training

Context transition trials were similar to preliminary training, with 16 odors presented as new in each arena and one S+ and one S-odor presented in each trial. In the two-context transition task, rats completed the first eight trials in the first arena, then all 16 trials in the second arena, and then returned to the first arena to complete the final eight trials of the list (Context A → Context B → Context A). So, when the rat was in the second arena, all of the odors were new to the context, even though half of them had been presented earlier in the session in the other context. Then, when the rat returned to the first arena, all of the odors had been presented in the session, but half were new to the context, and half were old in that context. In the three-context transition task shown in Figure 1, the rats completed the first eight trials in the first arena, then the first eight trials in the second arena, then returned to the first arena to complete the final eight trials in that context, then returned to the second arena to complete the final eight trials (Context A → Context B → Context A → Context B) (Figure 1).

**Figure 1.**
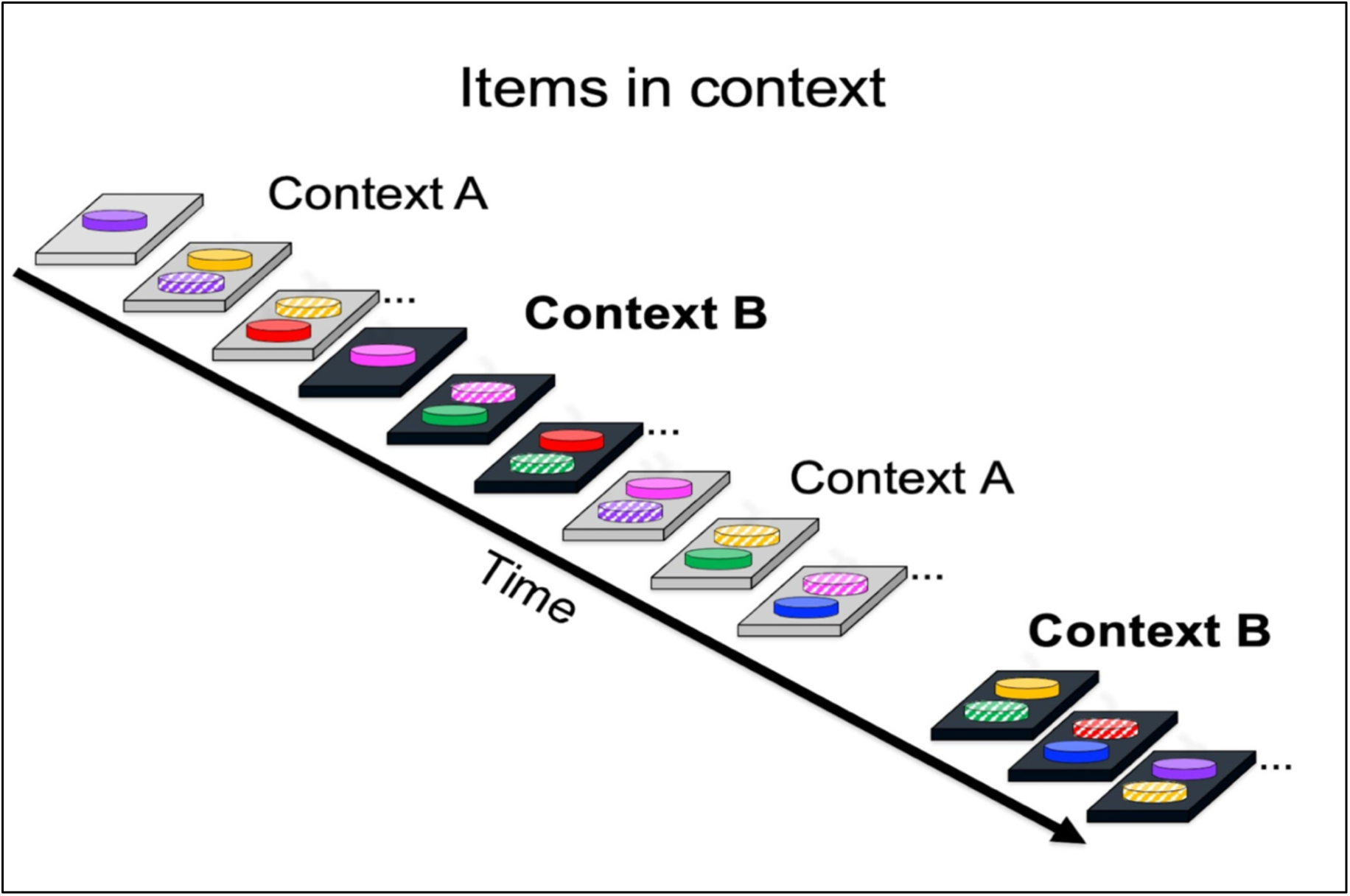
Schematic of the Item-in-context episodic memory task. This shows a trial divided across four segments. Rats completed eight trials in the first arena, then eight trials in the second arena, returned to the first arena to complete eight trials, and then returned to the second arena to complete the final eight trials. Light and dark grey rectangles depict arenas. Odors are represented by colored disks; the first presentation of an item in an arena is depicted by a solid color, whereas the same color with stripes depicts a subsequent presentation of an odor in the same arena. Selection of a new item (first presentation in the arena) is rewarded, whereas selection of an odor that has been presented previously in the same arena is not rewarded. Note that each odor is presented as new (i.e., rewarded) in each arena.

### Dissociating Episodic Memory from Familiarity

The list of odors was presented in two contexts, Context A and Context B. This is a challenging task because the odors from Context A will be used in Context B, but they will be considered “new” in each context using a random order of odors (Figure 1). Rats may remember all the items and the context in which they occurred using episodic memory (Panoz-Brown et al., 2016). However, rats may use familiarity cues instead of episodic memory to choose the correct answer for some sequences of odors. Figure 2A shows an example of how episodic memory and non-episodic memory (i.e., familiarity) can be confounded, with red representing the strawberry odor and blue representing blueberry. First, blueberry is presented as “new” in context A, and the rat is rewarded when choosing blueberry. Then, after transitioning to context B, the rat is presented with strawberry as the “new” odor. At this stage, blueberry has not yet been presented in context B. After returning to context A, strawberry is presented as the “new” odor in a later trial. Finally, the rat returns to context B and is presented with a choice between strawberry and blueberry.

**Figure 2.**
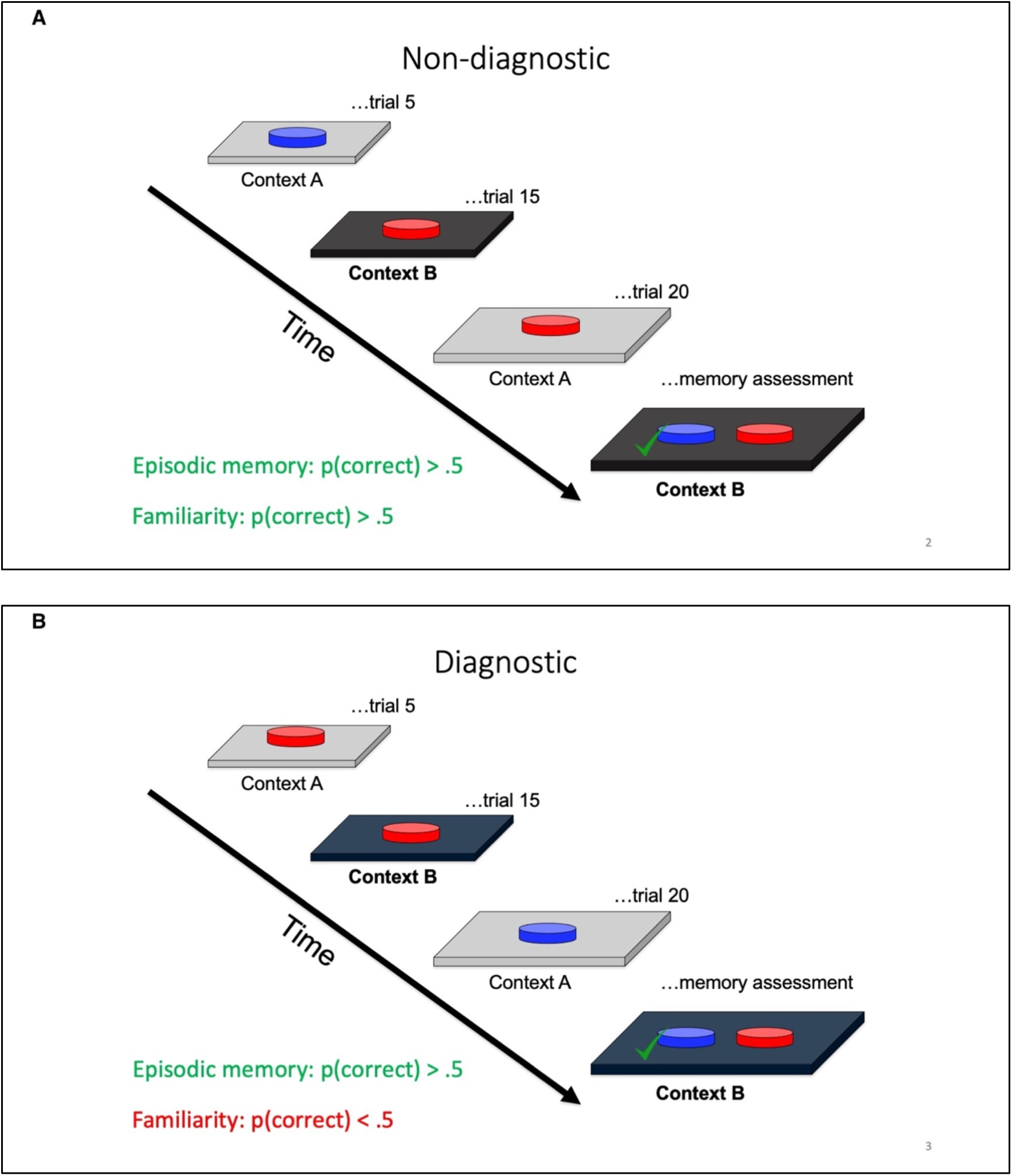
Familiarity and episodic memory are confounded in (A). A small change in the order of items unconfounds these two alternatives, as shown in (B). The memory assessment in (B) dissociates episodic memory (above chance) from judgments of relative familiarity (below chance). The presence of additional odors (not shown) is identified by “…” in the schematic. The schematic focuses on food-rewarded (S+) items (denoted by “√”) by omitting comparison with non-rewarded (S−) items prior to the memory assessment. Trials depicted in (A) and (B) are randomly intermingled throughout daily testing.

In this instance, blueberry is the correct choice, as the rat has not yet encountered blueberry in context B. If the rat relies on episodic memory to remember the items and contexts in which they were presented, it will choose blueberry. However, the rat may also choose blueberry if it relies on familiarity cues. In this case, note that blueberry was presented before strawberry. If the rat relies on the “avoid familiar item” rule, strawberry would seem more familiar because strawberry was presented as “new” more recently than blueberry. It would avoid strawberry and select blueberry, which is the correct choice. This is a confound, and sequences like this one are considered *non-diagnostic* for episodic memory because they can be answered correctly using either episodic memory or familiarity. As described next, a small change in the sequence dissociates episodic and non-episodic memory.

Figure 2B shows an example of how episodic memory and non-episodic memory (i.e., familiarity) can be dissociated. First, strawberry is presented as “new” in context A, and the rat is rewarded when choosing strawberry. Then, after transitioning to context B, the rat is presented with strawberry as the “new” odor. Blueberry has not yet been presented in context B at this stage. After returning to context A, blueberry is presented as the “new” odor. Finally, the rat returns to context B and is presented with a choice between strawberry and blueberry. In this scenario, blueberry is the correct answer because it was not yet presented in context B, and a rat using episodic memory (i.e., memory of all the items and the context in which they occurred) would choose blueberry.

If a rat relied on familiarity cues, it would avoid blueberry and choose strawberry in Figure 2B. Blueberry was presented as “new” more recently than strawberry, so blueberry is more familiar. By relying on familiarity in this scenario, the rat would choose strawberry, which is the incorrect choice. So, the only way to make the correct choice is by relying on episodic memory for all the items and the contexts in which they occurred. Notably, a rat relying on episodic memory will choose the correct item in context (above chance performance), whereas a rat relying on familiarity will choose the incorrect item (below chance) in Figure 2B. Therefore, trials with the sequence shown in Figure 2B are referred to as *diagnostic* for episodic memory. This dissociation of episodic memory from familiarity is an important aspect of the item-in-context task. The arrangement of odors was randomly selected in each arena, so the number of diagnostic and non-diagnostic trials varied daily.

### Mild Traumatic Brain Injury Animal Model

Rats were required to have at least 75% accuracy in the last five sessions of three context transition training before sham/injury manipulation. Once baseline data of three context transitions were obtained, an mTBI was induced with a modified version of the Wayne State University weight drop (WD) model. Our modification to the WD model was the introduction of an electromagnet to release/initiate the weight drop. Rats were anesthetized using sodium isoflurane. The rat was then placed prone on a slotted aluminum foil sheet taped to a U-shaped plexiglass container. Anesthesia was maintained using a nose cone. A foam pad was placed at the bottom of the plexiglass container. The rat’s head was positioned under a clear plexiglass guide tube where an electromagnet (Model X) released a 450 g cylindrical steel weight from a 1 m height. The rat then breaks through the slotted aluminum sheet and does a 180° rotation to land on its back on the foam pad. The weight was attached to a high-tension fishing line to avoid a second impact. The nose cone was removed before the weight drop. The rat was removed from the foam pad and returned to its home cage. This procedure produces an acceleration/deceleration injury that is typical of a mild head injury (Masse et al., 2019). Sham induction followed the same procedure except that the weight was not dropped. A randomized block design was used to assign rats to either the WD or the sham group based on baseline performance (5 sham, 6 WD) using diagnostic trial accuracy in the last five sessions before injury. Rats rested for 24 hours following sham/WD manipulation. Rats resumed daily behavioral testing sessions 24 hours after sham or weight drop manipulation with three context transitions for eight days. The equipment used in WDM is shown in Figure 3.

**Figure 3.**
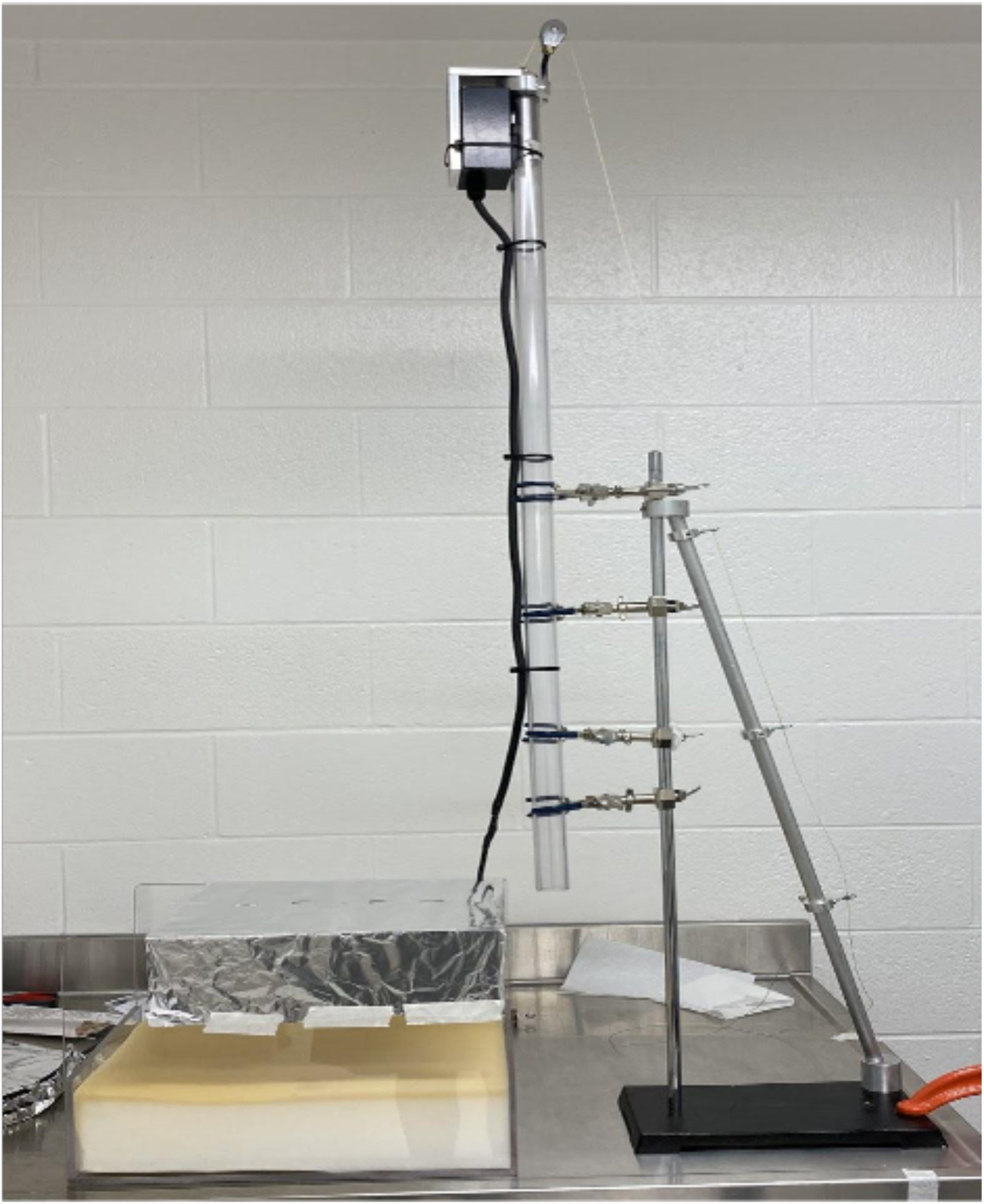
Image of Wayne State University Closed Head Weight Drop Model

### No Food Probes

A series of unbaited probes were conducted to test for possible odor detection of the pellets in the baited cup. Unbaited trials were conducted during the last session of three context transitions. For each rat, four probe trials were conducted in random order, with the constraints that at least one probe trial occurred in each context and the exception that the first trial in each context was not a probe trial. The new S+ and the old comparison stimulus were both unbaited in probe trials. Next, the experimenter manually delivered the chocolate pellet to the cup immediately following the rat’s selection of the S+ lid.

### Immunohistochemistry

For the rats used in the histological study, brains were collected from sham and WD rats one day (i.e. 24 hours post weight drop), four days, and eight days after injury. A four-day sham and a non-manipulation control group were also included (n = 6 per group). At the time of sacrifice, rats were deeply anesthetized with urethane via intraperitoneal injection, perfused with 4% paraformaldehyde (PFA) fixative, postfixed for 24 hours in PFA, and cryoprotected in 30% sucrose for three days. Brains were sliced at 40 microns from bregma −3.30 to −4.30 to obtain hippocampal slices. Slices were stained with Iba1, a marker for microglia, and GFAP, a marker for astrocytes. Free-floating sections were washed with 0.1M PBS and 0.3% TritonX100 PBS. Sections were blocked in 4% normal goat serum and incubated overnight at 4 °C with Iba1 and GFAP. The next day, slices were washed with 0.3% TritonX100 PBS and incubated with AlexaFlour 488, AlexaFlour 568, and DAPI for 2 hours. After incubation, slices were slide-mounted and left to dry overnight. The following day, slides were cover slipped using Prolong Diamond, and left to dry overnight. Slides were later imaged on a fluorescence microscope at 20x. The images were stitched together using the Lecia LAS X software.

### Morphology Analysis

An automated pipeline for measuring astrocyte and microglia morphology was developed in MATLAB and ImageJ, and the code is freely available at https://github.com/Nicholas-Port. The pipeline for segmenting the area of the cells is as follows: 1) A Gaussian blurred image is subtracted from the primary image to remove local changes in the background. 2) A Butterworth filter is applied to smooth the image. 3) The image is thresholded at the 95^th^ percentile to only examine bright pixels. 4) The image is sharpened with the localcontrast, locallapilt (local LapLacian filter) and imsharpen (unsharp mask) functions. 5) The edges of the image are detected with the Canny edge detection algorithm, and the image is converted to binary. 6) The binary images are then dilated, area-closed, filled, and eroded twice. 7) Objects smaller than 16 microns or larger than 1949 microns in area were removed. 8) Finally, the image is dilated three more times. For astrocytes or microglia to be accepted in the final analysis, the center of a DAPI segmented nucleus needs to be contained within the border of the segmented glial cell. Figure 4 shows examples of hippocampal tissue with astrocytes in green and DAPI-stained nuclei in blue. Superimposed on the image is the automated segmented morphology of the cells. Magenta outlines are accepted cells containing nuclei, and red outlines are unaccepted cells that do not contain a nucleus.

**Figure 4.**
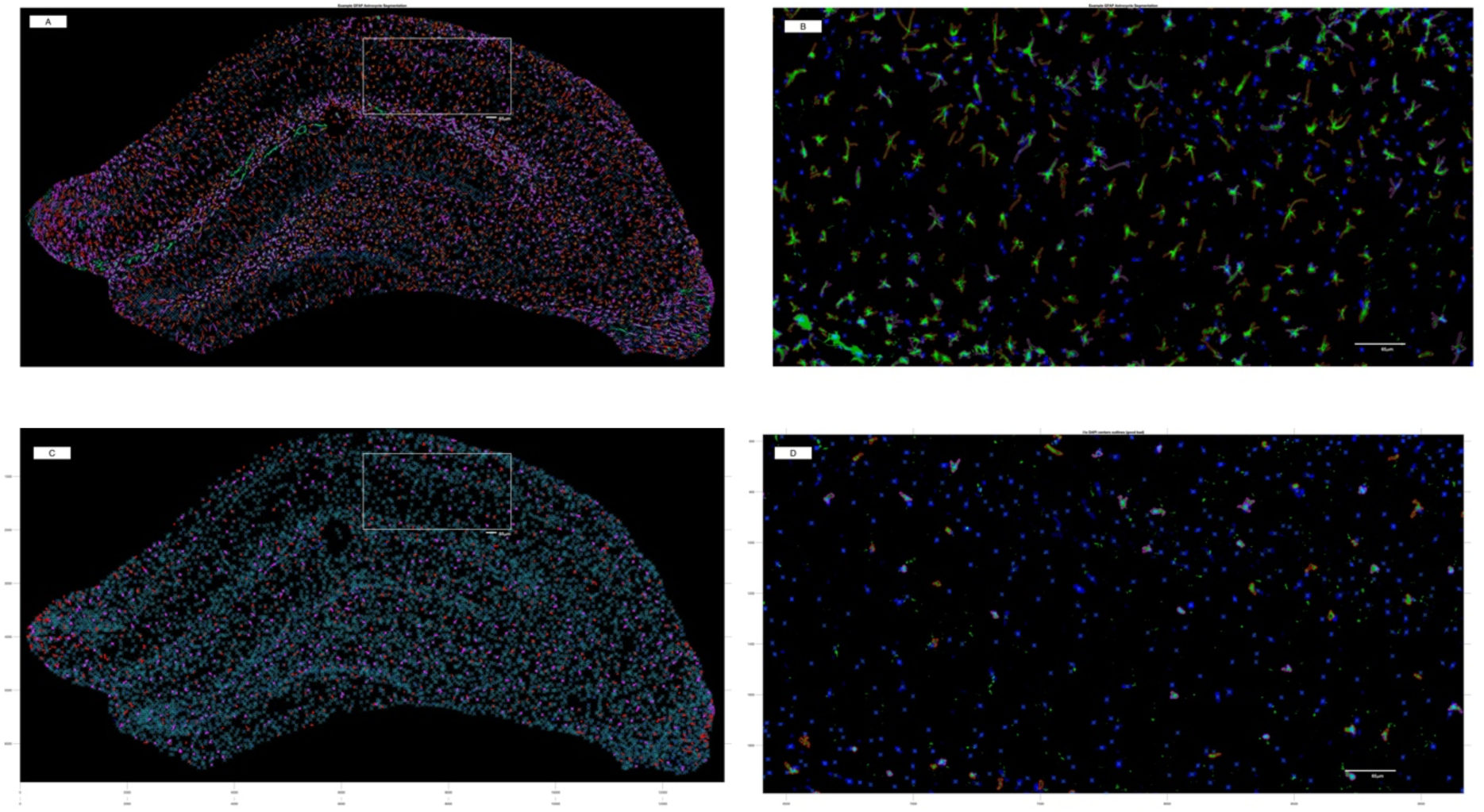
Representative images of the hippocampus and the segmentation analysis for Astrocytes (A-B) and Microglia (C-D). Positively stained astrocytes and microglia are highlighted in green in the zoomed-in images (B & D). In order for astrocytes or microglia to be accepted in the final analysis, the center of a DAPI-stained segmented nucleus (blue) must be contained within the border of the segmented glial cell. Magenta outlines are accepted cells containing nuclei, and red outlines are unaccepted cells that do not contain a nucleus (B & D).

A two-sample Kolmogorov-Smirnov test was initially used to determine whether the number or area of iba-1 or GFAP-expressing cells was observed between the naïve and sham groups. Identical conclusions were obtained using a two-tailed t-test. The sham-operated and naïve groups were subsequently pooled into a single control group for each dependent measure. A One-way ANOVA was subsequently performed to determine whether differences in either the number or area of iba-1 or GFAP-expressing cells were observed as a function of time post-weight drop versus the pooled control group. A Dunnett’s post hoc test was used to compare the number or area of iba-1 or GFAP-expressing cells at each time point to the pooled control group. Bonferroni’s multiple comparison tests were also used to compare these parameters derived from the group euthanized at 4 days post weight drop (i.e., the time point associated with an impairment of diagnostic episodic memory) with other post-weight drop time points. Bonferroni’s multiple comparison test used the same Mean Square Error term from the overall ANOVA, which consequently limits the possibility that a Type 2 error would be made by focusing post hoc comparisons on the critical comparisons of interest rather than on all possible comparisons. *P < 0.05* was considered statistically significant (GraphPad Prism software, San Diego, CA).

## RESULTS

### Baseline Performance

Baseline performance consisted of the mean accuracy of the last five days before sham/injury manipulation. The WD group (n= 6) had an average accuracy of 92.5% (SEM = 3.0%), and the sham group (n= 5) had an average accuracy of 92.1% (SEM = 3.4%) in the diagnostic trials. The WD group had an average accuracy of 93% (SEM = 2.7%), and the sham group had an average accuracy of 92% (SEM = 5.2%) in the non-diagnostic trials. The behavioral task results are summarized in Figure 5.

**Figure 5.**
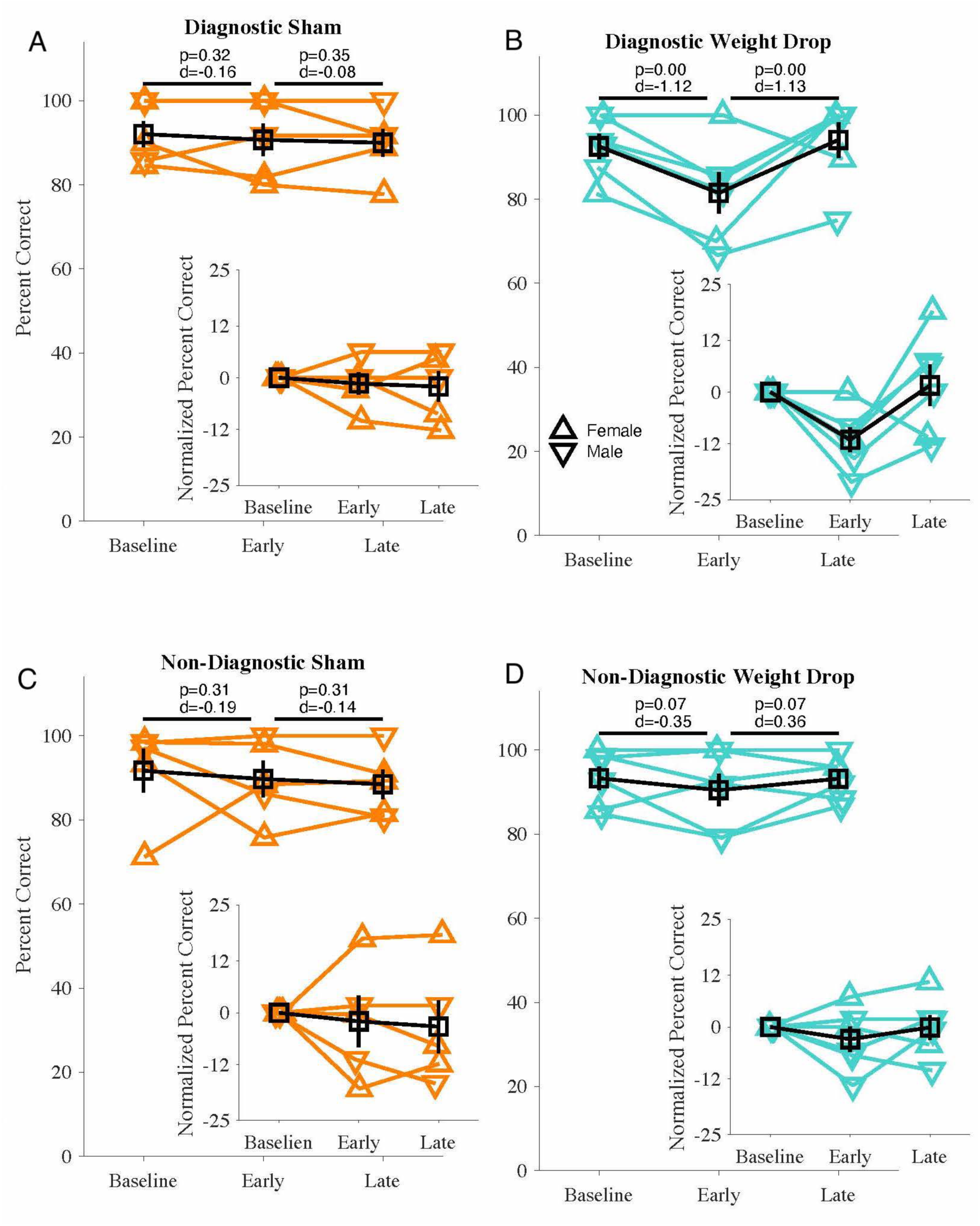
Percent correct and percent change from baseline on trials that are diagnostic of episodic memory (referred to as Diagnostic) and trials that are not diagnostic of episodic memory (referred to as non-diagnostic) in the sham (orange) and weight drop (orange) groups during the early and late time points. Mean data are shown as mean ±1 SEM. 2

### Post-Manipulation Performance

The post-manipulation data was divided into two timepoints: *early*, the first four days of post-manipulation testing, and *late*, the final four days of post-manipulation testing. The change in post-manipulation performance relative to baseline performance was assessed using a separate within-subjects t-test for *diagnostic* and *non-diagnostic* trials in the sham and WD groups (Figure 5). Figure 5A describes the performance of the sham (uninjured) group in the diagnostic trials. There were no significant changes in performance from baseline to the early timepoint (t(4) = −0.4, p > 0.05, d = 0.16), and from the early to the late timepoint (t(4) = −0.4, p > 0.05, d = 0.08). No significant difference was documented in the normalized baseline change data. Figure 5B describes the performance of the weight drop group in the diagnostic trials (trials that specifically test episodic memory). Rats’ performance significantly declined from baseline to the early timepoint (t(5) = −4.0, p < 0.01, d = 1.12). Performance significantly increased from the early to late timepoints (t(5) = −4.0, p < 0.01, d = 1.13). This is observed in the raw data and the normalized data insert, which shows the percent change from baseline for each timepoint. Figure 5C describes the performance of the sham group in the nondiagnostic trials. There were no significant changes in performance from baseline to the early timepoint (t(4) = −0.53, p > 0.05, d = 0.19), and from the early to the late timepoint (t(4) = −0.53, p > 0.05, d = 0.14). No significant difference was documented in the normalized baseline change data. Figure 5D describes the performance of the weight drop group in the nondiagnostic trials (trials that do not specifically test episodic memory). Performance did not significantly change from baseline to the early timepoint despite a slight decline in accuracy (t(5) = −0.96, p > 0.05, d = 0.35). This is observed in the normalized data, which shows a slight decline from the baseline at the early timepoint. No significant change occurred from the early to late timepoint despite a slight increase in accuracy (t(5) = −0.96, p > 0.05, d = 0.36).

### No Food Probes

If the high accuracy in baited trials is due to an ability to detect the baited food pellet, then in the absence of the pellet, it is expected that the animals would perform at the level of chance (50% correct). For no-bait trials, the two groups performed significantly above chance. The WD had an accuracy above chance (t(5) = 6.3, p < 0.01), and the sham group had an accuracy also above chance (t(4) = 6.5, p < 0.01), indicating that above chance accuracy was not due to the detection of the food pellet.

### Immunohistochemistry

A time course study was performed to ascertain the impact of a single weight drop on histological measures of glial cells expressing histological markers iba1 and GFAP (Fig. 6A-H). No statistically significant differences were observed between sham-operated and naïve groups in either the number of cells expressing iba-1 (Fig. 6A) or the area of iba1-expressing cells (Fig. 6C). Similarly, no statistically significant differences were observed between sham-operated and naïve group in either the number of cells expressing GFAP (Fig. 6E) or the area of GFAP-expressing cells (fig. 6D). Consequently, the sham-operated and naïve control groups were pooled into a single experimental control group for further statistical analysis.

**Figure 6.**
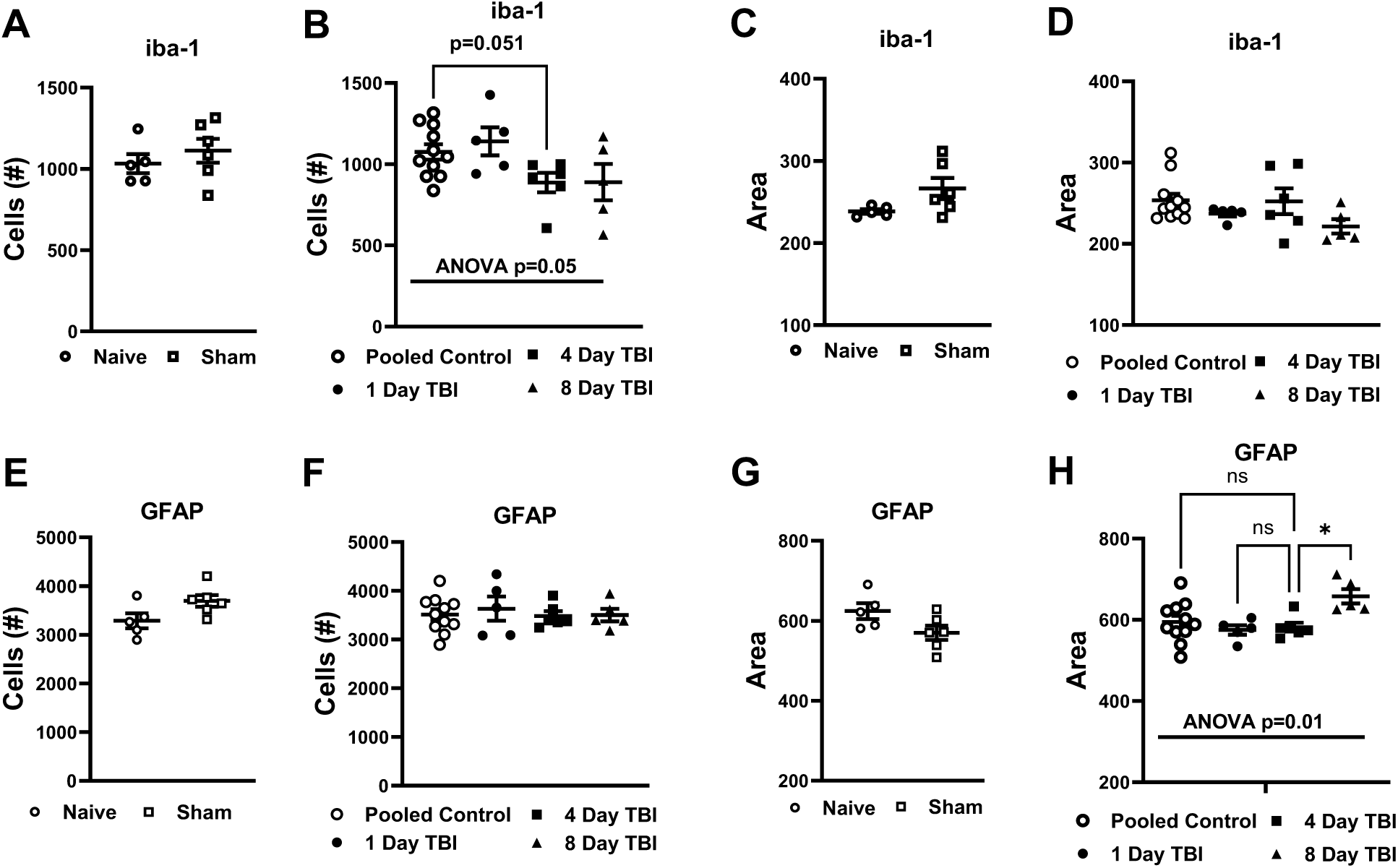
Histological measures for IBA-1 (microglia) (A-D) and GFAP (astrocytes) (E-H). Cell counts (A-B, E-F) and area (C-D, G-H) were used to analyze neurological damage in the hippocampus. Data are mean ± SEM. A one-way ANOVA followed by a Dunnett’s Test was performed to test the null hypothesis of whether there is an effect of group on each measure. * p < 0.05

One-way ANOVA revealed an overall trend (p=0.05) towards group difference in the number of iba-1 expressing cells in the hippocampus as a function of time post-weight drop. Multiple comparison tests revealed that there were fewer iba-1 expressing cells in the 4-day post-concussion injury group compared to the pooled control group. Dunnett’s multiple comparison post hoc test revealed a trend toward fewer iba-1 expressing cells in rats perfused at four days post weight drop compared to the pooled control (Figure 6B, p = 0.051). No reliable differences were observed between the 4-day post TBI compared to other time points (Figure 6B). One-way ANOVA failed to reveal reliable differences in the area of iba-1 expressing cells between groups (Fig. 6C, D). Similarly, no reliable differences in the number of GFAP-expressing cells were observed as a function of time post-TBI in the number of GFAP expressing cells (Fig. 6F). By contrast, one-way ANOVA revealed significant differences in the area of GAP-expressing cells as a function of time-post TBI. Bonferroni’s multiple comparison test revealed that the number of GFAP-expressing cells was greater in the 8-day post TBI group compared to the 4-day post TBI group (Fig. 6H). No other differences between groups were detected.

## DISCUSSION

This study was divided into two parts. The first was to determine if a mTBI impairs episodic memory in rats. We used the Wayne State University closed-head weight drop model to produce an mTBI. We used the Item-in-context task, which dissociates episodic memory from non-episodic solutions, to evaluate human-like episodic memory in rats (Panoz-Brown et al., 2016). In the trials that are diagnostic for episodic memory, we saw an immediate decline in performance in the injury group, which subsequently returned to baseline. We did not observe a decrease in performance in the sham group. In trials that are non-diagnostic for episodic memory, we did not observe any changes in performance.

The decline in diagnostic trials with the preservation of performance on non-diagnostic trials suggests the Wayne State University injury model specifically affected episodic-like memory in rodents while sparing the other aspects of memory. This could be because episodic memory is hippocampal dependent, whereas non-episodic memory is not hippocampal dependent (Aggleton et al., 2005; Yonelinas, Otten, Shaw, & Rugg, 2005). During our initial observation of the rats’ day-by-day behavioral performance, we noticed the WD group’s peak deficit immediately after injury but subsequently returned to baseline. Due to the recovery of performance, we split the data into two timepoints. Days 1-4 were *early*, and 5-8 were *late*. This allowed us to document both an injury and recovery effect.

Since the accepted definition of episodic memory is the ability to remember specific events and the context in which they occur (Crystal, 2009; Eichenbaum, 2001; Tulving, 2002), the Item-in-context task (Panoz-Brown et al., 2016) was an appropriate approach to examine episodic memory function in rats in this study. This adds to the research showing that rats can exhibit human-like episodic memory. Furthermore, the disruption of episodic memory function adds to the validity of the WDM. This WDM interfered with a key aspect of episodic memory without the limitations commonly seen with other models of mTBI (Masse et al., 2019; Qin et al., 2018).

Additionally, we can produce human-like behavioral deficits by using a model that replicates the biomechanics of human injury (Hiskens et al., 2019; Masse et al., 2019; Romeu-Mejia et al., 2019). Therefore, the observed impairment in episodic memory using this WDM aligns with our primary hypothesis.

Based on the behavioral data, we evaluated the timecourse of changes in the number and area of iba-1 and GFAP expressing cells in the hippocampus following a single weight drop. The weight drop was sufficient to produce deficits in our diagnostic episodic memory assessment at 4-days post TBI. We focused on the expression of GFAP-expressing astrocytes and iba-1 expressing microglia/macrophages since these markers are commonly used cellular markers of neuroinflammation that have been linked to concussion injury (Lafrenaye et al., 2015; Sofroniew & Vinters, 2010; Zhou et al., 2020). Since the peak behavioral deficit occurred four days post-injury and returned to baseline eight days post-injury, it was important to monitor the expression of astrocytes and microglia across a range of timepoints.

We observed a significant increase in the area of GFAP-expressing cells from four days to eight days post-injury. GFAP expression and area may increase following more transient changes in microglial activation. Our studies suggest that, despite episodic memory performance returning to baseline by eight days post-injury, the area of astrocytes continued to increase following the injury. More work is necessary to ascertain whether this change in area of GFAP-expressing cells reflects an adaptive change in response to other changes in the cellular milieu. We did not observe a similar increase in area of iba-1 expressing activated microglia. One could speculate the time course of changes in cell size or expression is different for these two cell populations (Grovola et al., 2021; Zhou et al., 2020); changes in the number or shape of activated microglia are largely believed to precede changes in astrocytes. Whereas microglia are the primary phagocytes that clear cellular debris in the central nervous system, astrocytes, which can exhibit pro-inflammatory gene expression profiles, can engulf microglial debris (Konishi, 2020) as a compensatory mechanism to reinstate a healthy CNS. It is also possible that changes in microglia and astrocytes in different subregions of the hippocampus could be masked by our global measurements of changes in the number or area of iba-1 or GFAP expressing cells in the entire hippocampus, rather than more discrete subregions of the hippocampus (e.g., dentate gyrus, CA1) or other gray or white matter regions. It is also likely that repeated TBI would produce more pronounced changes in the expression of these markers compared to the single weight drop employed here. Consistent with this hypothesis, repeated close head injury induced TBI increased expression of iba-1 in CA1, dentate gyrus (DG), corpus callosum, and ventral tegmental area/substantia nigra pars compacta relative to sham or 1-day post single concussion-injury induced TBI (Broussard et al., 2018). In this latter study, performed in mice, effects of a single closed-head injury-induced TBI on GFAP expression did not differ reliably from sham animals in any brain region, whereas robust elevations in GFAP were observed following repeated TBI. It is also possible that differences in numbers or area of iba-1 or GFAP expressing cells are observed between male and female rats, limiting our ability to detect group differences across time. We did not evaluate sex differences in our histological markers due to small sample sizes and limitations in the throughput of episodic memory function, but future studies could explore this possibility, particularly focusing on early timepoints (i.e., less than 1-day post TBI).

We observed a trend towards a decrease in microglia cell counts across days, particularly on day four post-injury, whereas the number of astrocytes did not change across days. By contrast, the area of GFAP-expressing cells increased on day 8 relative to day 4 post-TBI. Our studies raise the possibility that the increased area of GFAP-expressing cells may be related to astrocyte engulfment of activated microglia, as described above (Konishi, et al., 2020; Konishi et al., 2022) Hence, the area of GFAP-expressing cells was increased subsequent to our observation of decreases in the number of iba-1-expressing cells. Following a single weight drop, it is possible that an increase in the number of iba-1 expressing cells could be transiently observed before the first 1-day post-TBI timepoint.

Consequently, elevation in the number of iba-1 expressing cells may have peaked prior to the 1-day post-TBI perfusion time point. Changes in astrocytes and microglia may document hippocampal dysfunction. Consistent with this hypothesis, repeated concussion injury has been linked to the expression of astrocytes and microglia. One role of the hippocampus is to bind spatial and non-spatial information, which creates context-dependent memories. Context-dependent memories are crucial for episodic memory function (Lee & Jung, 2017). Episodic memory has been shown to be hippocampal-dependent (Tulving & Markowitsch, 1998). Thus, damage to the hippocampus may impede episodic memory function (Bai et al., 2009; La et al., 2019; Panoz-Brown et al., 2018). Hence, the observed hippocampal damage caused by the Wayne State WDM may have produced episodic memory deficits, which aligns with our secondary hypothesis. We did not explore possible changes in white matter, which have been increasingly detected following repeated TBI in other models.

This study was able to produce human-like episodic memory impairment in rats without some of the limitations of other injury models (fixed head, skull fractures, high mortality rate, etc.). Yet, some limitations are noted. This study used male and female rats in the behavioral analysis and the histology timecourse study. Due to the small number of males and females in each group, we did not evaluate sex differences in this study. As mentioned above, we conducted a histology timecourse study on a separate group of mixed-sex (male and females were combined) rats. It is possible that the rats who performed the behavioral portion of this experiment may have exhibited a different rate of astrocyte and microglia expression due to the added responsibility of doing the behavioral task.

The time course study analyzed the entire hippocampus. Previous studies have shown that lesions in parts of the hippocampus can interfere with the ability to form associations between contexts and odors. Future studies may examine if there are region-specific changes in astrocytes and microglia (CA1, CA3, DG, etc.) over several days. In the human literature, people can experience multiple mTBI throughout their lives (Dewitt et al., 2013). Future studies should investigate episodic memory impairment following multiple injuries as a model of repetitive mTBI using the Wayne State model. Our study specifically focused on a single concussion injury to better enable comparisons with a single concussion injury in our concurrent study assessing episodic memory impairment following acute concussion injury in human athletes.

## CONCLUSIONS

Mild traumatic brain injury (concussion) affects millions of people every year. Memory impairment is one of the most common symptoms following mTBI. The Wayne State University WDM is a more human-like injury model than the other standard animal models by modeling human injury biomechanics. Our study is the first to show an episodic memory deficit using an animal model of mTBI. These findings validate our injury model and show we can model human-like episodic memory impairment in rats.

## ACKNOWLEDGMENTS

This work was supported by NSF/BCS-1946039 to JDC, and GDN was supported by NIMH 5 T32 MH103213.

